# Disturbed repolarisation-relaxation coupling during acute ischaemia permits systolic mechano-arrhythmogenesis

**DOI:** 10.1101/2024.12.19.629548

**Authors:** Breanne A. Cameron, Peter A. Baumeister, Tarek Lawen, Sara A. Rafferty, Behzad Taeb, Matthew R. Stoyek, Joachim Greiner, Ilija Uzelac, Flavio H. Fenton, Rémi Peyronnet, Peter Kohl, T Alexander Quinn

**Author notes:** CORRESPONDING AUTHOR: Breanne A. Cameron, Institute for Experimental Cardiovascular Medicine, Universitätsklinikum Freiburg, Universitäts-Herzzentrum, Elsässer Straße 2Q, 79110, Freiburg, Germany.

## Abstract

**Background:** The heart’s mechanical state feeds back to its electrical activity, potentially contributing to arrhythmias. ‘Mechano-arrhythmogenesis’ has been mechanistically explained during electrical diastole, when cardiomyocytes are at their resting membrane potential. During electrical systole, cardiomyocytes are refractory right from the onset of depolarisation, while during repolarisation they appear to be protected from mechano-arrhythmogenesis by near-simultaneous restoration of resting membrane potential and cytosolic calcium concentration ([Ca^2+^]_i_): repolarisation-relaxation coupling (RRC). Yet, systolic mechano-arrhythmogenesis has been reported in ischaemic myocardium, with unclear underlying mechanisms. We hypothesise that ischaemia-induced alteration of RRC gives rise to a vulnerable period for mechano-arrhythmogenesis.

**Methods:** Acute left-ventricular (LV) regional ischaemia was induced by coronary artery ligation in Langendorff-perfused rabbit hearts, with mechanical load controlled by an intraventricular balloon. Mechanical activity was assessed by echocardiography and arrhythmia incidence by electrocardiogram. Single LV cardiomyocytes were exposed to simulated ischaemia or pinacidil (ATP-sensitive potassium channel opener). Stretch was applied in diastole or late systole using carbon fibres. Stretch characteristics and arrhythmia incidence were assessed by sarcomere length measurement. In both models, RRC was assessed by simultaneous voltage-[Ca^2+^]_i_ fluorescence imaging and mechano-arrhythmogenesis mechanisms were pharmacologically tested.

**Results:** In whole heart, acute regional ischaemia leads to systolic stretch and disturbed RRC at the ischaemic border. These electro-mechanical changes were associated with waves of arrhythmias, which were reduced by mechanical unloading, electro-mechanical uncoupling, or buffering of [Ca^2+^]_i_. In LV cardiomyocytes, physiological RRC is associated with a low incidence of systolic mechano-arrhythmogenesis, while a vulnerable period emerged by prolonged RRC during ischaemia. The increase in systolic mechano-arrhythmogenesis was reduced by restoring RRC, chelating [Ca^2+^]_i_, blocking mechano-sensitive transient receptor potential kinase ankyrin 1 channels (TRPA1), or buffering reactive oxygen species (ROS) levels.

**Conclusion:** Prolonged RRC allows for systolic mechano-arrhythmogenesis in acute ischaemia, involving contributions of elevated [Ca^2+^]_i_, TRPA1 activity, and ROS, which represent potential anti-arrhythmic targets.

**GRAPHICAL ABSTRACT LEGEND:** **Role of disturbed repolarisation-relaxation coupling (RRC), transient receptor potential kinase ankyrin 1 (TRPA1) channels, cytosolic calcium concentration ([Ca^2+^]_i_), and reactive oxygen species (ROS) in ventricular systolic mechano-arrhythmogenesis.** Schematic of the proposed mechanisms underlying the TRPA1- and Ca^2+^-mediated increase in systolic mechano-arrhythmogenesis with disturbed RRC. AITC, Allyl isothiocyanate (TRPA1 channel activator); AP, action potential; BAPTA ([Ca^2+^]_i_ buffer); CaT, Ca^2+^ transient; DNT, dantrolene (ryanodine receptor stabiliser); DPI, diphenyleneiodonium (ROS production blocker); GLIB, glibenclamide (K_ATP_ channel blocker); HC-300031 (TRPA1 channel blocker); K_ATP_, ATP-sensitive potassium channel; NAC, N-acetyl-L-cysteine (ROS chelator); NCX, sodium-Ca^2+^ exchanger; PIN, pinacidil (K_ATP_ channel activator); ROS, reactive oxygen species; SI, simulated ischaemia; STP, streptomycin (non-specific mechano-sensitive ion channel blocker).

## INTRODUCTION

Feedback is an essential element of biological systems’ autoregulation, and fundamental to the adaptation of physiological activity to varying demands. A prime example is the heart – an electro-mechanical pump whose electrical excitation causes mechanical contraction of cardiomyocytes, involving a feedforward mechanism of electrically-triggered, calcium (Ca^2+^)- mediated Ca^2+^ release from intracellular stores (excitation-contraction coupling) and mechano-sensitive feedback by which the mechanical state of cardiomyocytes affects their electrical activity (mechano-electric coupling).^1^ Although mechano-electric coupling is important for maintaining and fine-tuning normal cardiac function,^2^ it can also contribute to arrhythmias by causing aberrant electrophysiological behaviour (‘mechano-arrhythmogenesis’),^3^ particularly in cardiac diseases that alter the finely-balanced physiological heterogeneity^4^ of passive and active mechanics of the heart.^5^ While the influence of mechano-electric coupling on the heart’s electrical activity is well established,^1^ the mechanisms and molecular identity of specific driver(s) underlying mechano-arrhythmogenesis – in particular during systole – are unclear.^6^

Mechano-arrhythmogenesis involving mechanical stimulation during ‘electrical diastole’ (the phase of the cardiac cycle in which cardiomyocytes are at their resting membrane potential [V_m_], and hence fully excitable) has been reliably explained. This has been shown to involve mechanically-driven excitation through cardiomyocyte depolarisation by mechano-sensitive ion channels and modulation of intracellular Ca^2+^ handling,^1^ including stretch-induced increases in sarcoplasmic reticulum Ca^2+^ release,^7^ or stretch-and-release related surges in free cytosolic Ca^2+^ concentration ([Ca^2+^]_i_) due to stretch- and release-effects on myofilament Ca^2+^ binding.^8^ On the other hand, under physiological conditions cardiomyocytes appear to be well-protected from mechano-arrhythmogenesis during ‘electrical systole’ (while the action potential [AP] is occurring). This includes both the early (cell excitation and AP plateau) and late phases (period during which V_m_ and [Ca^2+^]_i_ are being restored to resting values) of systole. This restoration involves progressive V_m_ repolarisation, which prevents further Ca^2+^ influx through voltage-gated channels, but also increases the availability of fast Na^+^ channels underlying recovery of cardiomyocytes from refractoriness, while simultaneously reducing [Ca^2+^]_i_ *via* sequestration back into intracellular stores and extrusion from the cell, ultimately leading to mechanical relaxation. This well-coordinated process – henceforth referred to as ‘repolarisation-relaxation coupling’ (RRC) – may account for the protection of cardiomyocytes from late systolic mechano-arrhythmogenesis (along with protection in early systole by electrical refractoriness that follows excitation). In pathological states, however, a dissociation of electro-mechanical recovery dynamics may limit this protection.^5^

One such pathology is acute regional ischaemia following coronary artery occlusion, which constitutes a major risk for sudden cardiac death.^9^ Pathological heterogeneity of myocardial mechanical properties and activity is thought to contribute to the induction of arrhythmias in acute ischemia, due to systolic stretch of less contractile myocardium at the border of the ischaemic region (often referred to as paradoxical segment lengthening).^10,11^ This is especially true for arrhythmias that occur ∼15-60 min after complete coronary artery occlusion (also known as ‘phase 1b’ ischaemia), as demonstrated in experimental^12^ and computational^13^ whole heart studies. The mechanisms of systolic mechano-arrhythmogenesis in acute ischemia are unexplored. We hypothesise that a disturbance of the normally well-coordinated systolic recovery of V_m_ and [Ca^2+^]_i_ during RRC is involved in mechano-arrhythmogenesis in ischaemic tissue^14–16^ during phase 1b of acute ischemia. This involves a pronounced shortening of AP duration (APD) by activation of ATP-sensitive potassium (K_ATP_) channels (arising as a consequence of reduced oxygen availability) and a less pronounced shortening of [Ca^2+^]_i_ transient duration (CaTD). As a result, RRC is delayed, giving rise to a vulnerable period for mechano-arrhythmogenesis in late systole, as high [Ca^2+^]_i_ favours arrhythmogenesis in progressively re-excitable cardiomyocytes.

We explored this possibility using rabbit Langendorff-perfused whole hearts and isolated left ventricle (LV) cardiomyocytes. Hearts were subjected to acute regional LV ischaemia, combined with: (i) control of LV mechanical loading using an intraventricular balloon; (ii) measurement of LV mechanical activity by speckle-tracking echocardiography and of arrythmia incidence by electrocardiography; (iii) assessment of RRC disturbance through dual V_m_-[Ca^2+^]_i_ optical mapping; and (iv) pharmacological interrogations of mechano-arrhythmogenesis mechanisms. Single LV cardiomyocytes were subjected to simulated ischaemia or pharmacological activation of K_ATP_ channels, combined with: (i) application of transient, timed stretch using carbon fibres; (ii) video-based measurement of sarcomere length dynamics; (ii) simultaneous fluorescence imaging of V_m_ and [Ca^2+^]_i_; and (iii) pharmacological interrogations. These experiments revealed prolonged RRC as a predictor of systolic mechano-arrhythmogenesis during acute ischaemia, identifying elevated [Ca^2+^]_i_, transient receptor potential kinase ankyrin 1 (TRPA1) activity, and reactive oxygen species (ROS) as underlying mechanisms.

## METHODS

### Data Availability

Key methodological information is provided below, with more comprehensive details in the Supplemental Material. The data contained in all figures is available on figshare (doi: TBD), and along with the computer code that support the findings of this study, are also available from the corresponding author upon reasonable request.

### Ethical Approval

Experiments used female New Zealand White rabbits (2.1 ± 0.2 kg, Charles River) and were conducted in accordance with the ethical guidelines of the Canadian Council on Animal Care and the German legislation for animal welfare, following protocols approved by the Dalhousie University Committee for Laboratory Animals and the local Institutional Animal Care and Use Committee at the University of Freiburg. Experimental methodologies followed the Guidelines for Assessment of Cardiac Electrophysiology and Arrhythmias in Small Animals^17^ and Guidelines for Experimental Models of Myocardial Ischemia and Infarction,^18^ with details reported following the Minimum Information about a Cardiac Electrophysiology Experiment (MICEE) standard.^19^

### Whole Heart Acute Regional Ischaemia Model

Hearts were perfused at 20 mL/min with a bicarbonate-buffered solution using a Langendorff-apparatus and instrumented with right atrial pacing electrodes and a LV balloon. The balloon was connected to a height-adjustable hydrostatic column for passive filling, with a diastolic pressure head of ∼5 mmHg. Afterload during ejection was controlled by an adjustable screw clamp around the balloon-column tubing, tightened such that maximum systolic pressure during ejection was ∼60 mmHg before ischemia or pharmacological interventions (Figure 1a). A suture was placed around the anterior branch of the left circumflex coronary artery,^20^ ∼1/3 of the distance between the LV base and apex, to obtain a ∼20% reduction in total coronary perfusion flow (when perfused with a constant pressure of 80 mmHg) and an ischaemic region of ∼40% of whole ventricular tissue volume upon ligation (which has been shown to result in the highest arrhythmia incidence in experimental models^21^; Figure S1). Hearts were placed in a custom-made imaging chamber, with bath solution maintained at 37°C and bubbled with 95% nitrogen and 5% carbon dioxide, to prevent trans-epicardial diffusion of oxygen into the ischaemic tissue.^22^ The chamber had a silicone imaging window for measurement of cardiac mechanical activity by two-dimensional echocardiography, and electrodes for contact-free pseudo-electrocardiogram acquisition (Figure 1a). Before induction of regional ischaemia, Langendorff perfusion was switched to a constant pressure of 80 mmHg (which resulted in a total coronary perfusion flow of ∼20 mL/min) and hearts were paced at 4 Hz, to ensure capture above the intrinsic heart rate. Hearts were excluded if the flow rate was < 15 mL/min. After a 10 min period for acclimatisation to constant pressure perfusion, the coronary artery was ligated for 60 min, during which time effects on mechanical activity and arrhythmia incidence were assessed (characterised as ‘ectopic excitation’, involving 1 or 2 consecutive ectopic QRS complexes and pressure waveforms out of sync with the atrial pacing stimuli, or ‘self-terminating arrhythmic activity’, involving 3 or more consecutive ectopic QRS complexes and pressure waveforms or disorganised activity in the case of non-sustained ventricular fibrillation; Figure 2a), followed by injection of 6 μm diameter fluorescent microspheres through the aortic cannula for characterisation of the ischaemic region by ‘negative staining’ (fluorescence-free tissue; Figure S1a).

**Figure 1.**
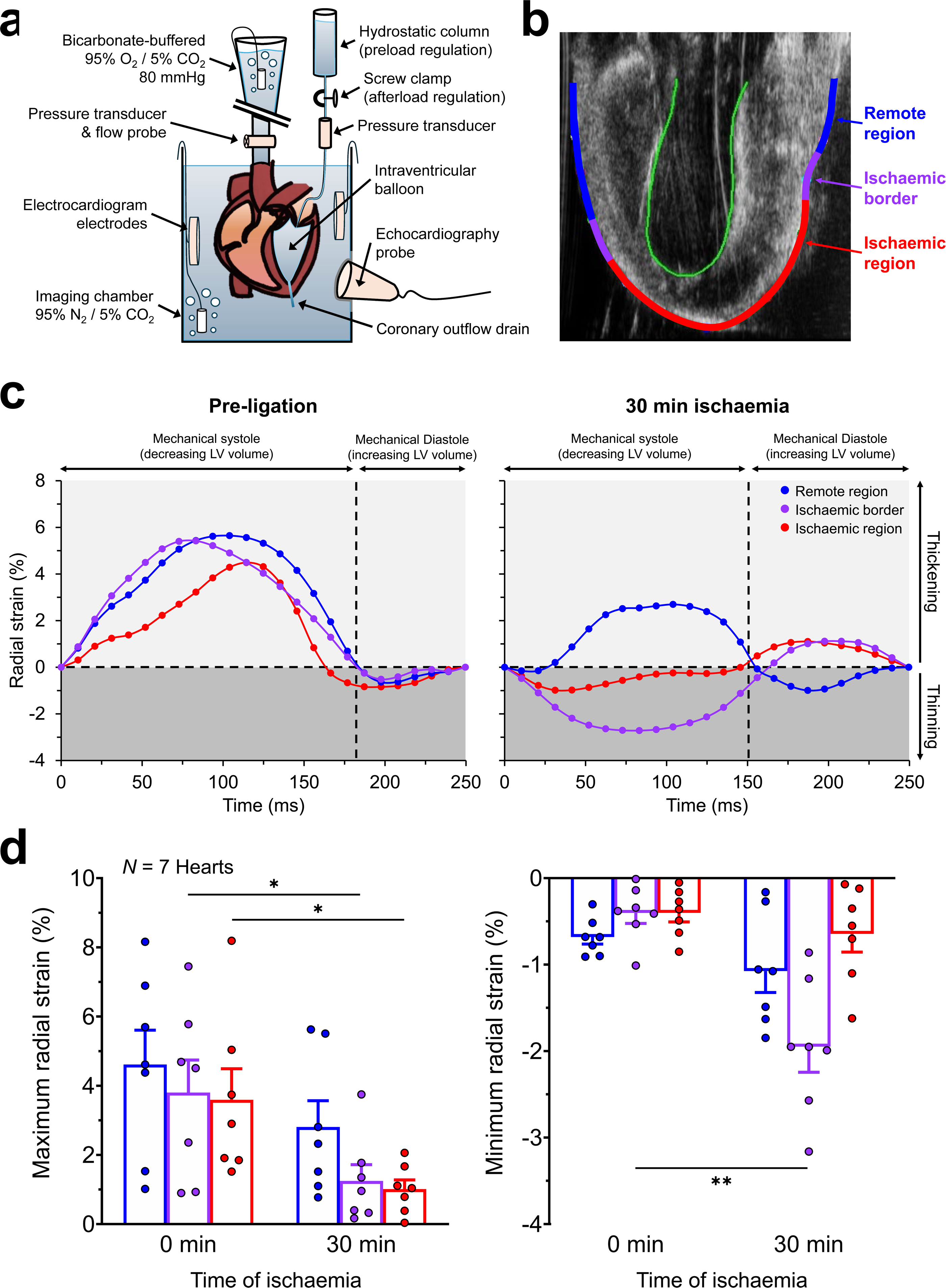
Altered left ventricular (LV) mechanical activity during acute regional ischaemia in rabbit whole hearts. **a,** Schematic illustration of the experimental setup for the whole heart regional ischaemia model. **b,** Representative two-dimensional echocardiographic image of the left ventricle showing the normally perfused remote tissue, the ischaemic border, and central ischaemic tissue regions. **c,** Example of measured radial strain over one heartbeat in each of the three regions before (left) and 30 min after (right) ligation of the anterior branch of the left circumflex coronary artery at ∼1/3 of the distance between the LV base and apex. **d,** Summary data on maximum (wall thickening, left) and minimum (wall thinning, right) radial strains before (0 min) and after (30 min) coronary artery ligation averaged over three beats in each heart, demonstrating an ischaemia-induced decrease in contractile function of ischaemic and border tissue, and stretch of ischaemic border tissue. Differences between pre- and 30 min post-ligation assessed by paired, two-tailed, Student’s *t*-tests. \**p* < 0.05, \*\**p* < 0.01.

**Figure 2.**
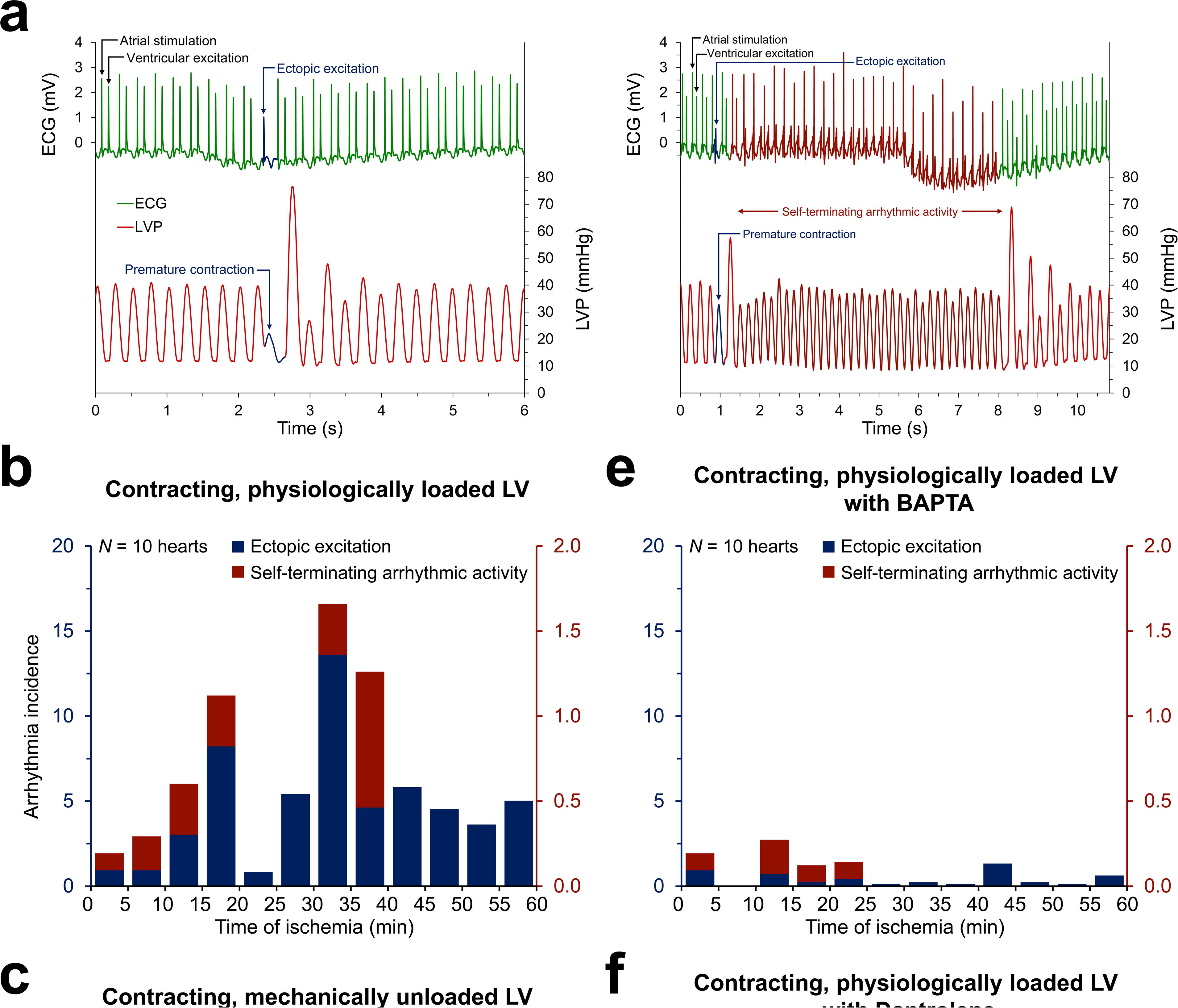
Arrhythmia incidence during acute regional ischaemia in rabbit whole hearts. **a,** Representative electrocardiogram (ECG) and left ventricular (LV) pressure recordings 30 min after coronary artery ligation, showing examples of ectopic excitation (left) and self-terminating arrhythmic activity (right). Incidence of ectopic excitation (blue, left y-axis) and self-terminating arrhythmic activity (red, right y-axis) over 60 min (binned into 5 min periods) after artery ligation. Experimental groups comprise hearts with a: **b,** contracting, physiologically loaded LV; **c,** contracting, mechanically unloaded LV; and **d,** non-contracting, physiologically loaded LV. In addition, results are shown for heart with a contracting, physiologically loaded LV that were treated with: **e,** BAPTA (to buffer cytosolic calcium) or **f,** dantrolene (DNT, to stabilise ryanodine receptors in their closed state). **g**, Average arrhythmia incidence in the 20 to 45 min window after artery ligation (phase 1b). Whether the LV of hearts in each group were physiologically loaded and/or contracting is indicated by the × marks below the x-axis labels. Differences between interventions and loaded hearts assessed by one-way ANOVA with Dunnet *post-hoc* tests. \**p* < 0.05, \*\**p* < 0.01. Note: for the self-terminating arrhythmic activity, there was one occurrence of non-sustained ventricular fibrillation in two of the contracting, mechanically unloaded hearts.

### Single LV Cardiomyocyte Model

For single cell experiments, hearts were Langendorff-perfused and LV cardiomyocytes were enzymatically isolated. Isolated LV cardiomyocytes were exposed to: (i) control solution (CTRL; containing, in mM: 142 NaCl, 4.7 KCl, 1 MgCl_2_, 1.8 CaCl_2_, 10 glucose, and 10 HEPES [Sigma-Aldrich]; pH adjusted to 7.40 ± 0.05 with NaOH; osmolality 300 ± 5 mOsm/L); (ii) a simulated ischaemia (SI) solution with a composition that aims to mimic extracellular fluid after 30 min of ischaemia^23,24^ (containing, in mM: 140 NaCl, 15 KCl, 1.8 CaCl_2_, 1 MgCl_2_, 10 HEPES, 1 NaCN to block oxidative phosphorylation, and 20 2-deoxyglucose to block anaerobic glycolysis [Sigma-Aldrich]; pH adjusted to 6.5 ± 0.05 with NaOH, osmolality 300 ± 5 mOsm/L); or (iii) CTRL solution containing pinacidil (PIN; 50 µM, to activate ATP-sensitive potassium [K_ATP_] channels).

### Carbon Fibre-Based Single Cell Stretch

Single cardiomyocytes, maintained at 35°C and paced at 1 Hz, were stretched using the carbon fibre method.^7,23^ Transient stretch, mimicking paradoxical segment lengthening of weakened myocardium during regional ischaemia,^25^ was applied to the cell (20 µm piezo-electric actuator displacement, applied and removed at a rate of 0.7 μm/ms, with a total stretch and release pulse duration of 110 ms; Figure 4a, b). Stretch was timed from the electrical stimulus to commence either during mid-diastole (600 ms delay), or in late systole during RRC (300 ms delay in CTRL, 150 ms during exposure to SI or PIN, and 210 ms during exposure to SI or PIN with glibenclamide, based on timings measured by simultaneous V_m_-Ca^2+^ imaging, described below). The intervention was repeated after 10 s at resting length for a total of four stretches (Figure 4b). This 4-stretch protocol was repeated with increased magnitudes of displacement (30 µm and 40 µm, with total stretch and release pulse durations of 140 ms or 170 ms) to generate a range of stretch-induced changes in sarcomere length within a cardiomyocyte (30 s between runs of 4, for a total of 12 stretches; Figure 4b). Sarcomere length and the piezo-electric actuator and carbon fibre tip positions were monitored and recorded at 240 Hz (Myocyte Contractility Recording System, IonOptix), to characterise cardiomyocyte mechanical properties and the mechanical effects of stretch, and to assess the incidence of mechano-arrhythmogenesis (characterised as ‘stretch-induced contraction’, involving 1 or 2 consecutive contractions after stretch out of sync with the electrical field stimulation or ‘self-sustained arrhythmic activity’, involving 3 or more successive contractions immediately after stretch that either spontaneously resolved or were terminated by an additional stretch; Figure 4c-f).

### Pharmacological Interventions

For whole heart experiments, mechanisms contributing to arrhythmogenesis during ischaemia were assessed by pre-treating hearts with pharmacological agents, including: the excitation-contraction uncoupler (±)-blebbistatin (to stop contractions; continuous reperfusion of 8 μM from a 10 mM stock in dimethyl-sulfoxide [Toronto Research Chemicals], with an initial 30 min incubation period); BAPTA-AM (to buffer [Ca^2+^]_i_; 0.1 µM from a 10 mM stock in dimethyl sulfoxide [Abcam] for 30 min [concentration determined in preliminary experiments by titrating to the maximum value that caused no change in developed LV pressure during contraction] followed by 30 min of BAPTA-AM free perfusion for de-esterification); and dantrolene (to stabilise ryanodine receptors; 1 µM from a 5 mM stock in dimethyl sulfoxide [Abcam] for 20 min).

For single cell experiments, mechanisms contributing to mechano-arrhythmogenesis were assessed by pre-treating cardiomyocytes with pharmacological agents, including: glibenclamide (to block K_ATP_ channels; 20 μM from a 10 mM stock in dimethyl sulfoxide [Abcam] for 15 min); BAPTA-AM (1 µM from a 10 mM stock in dimethyl sulfoxide [Abcam] for 20 min, with the concentration determined in preliminary experiments by titrating to the value that caused no more than a ∼10% decrease in percent sarcomere shortening during contraction); dantrolene (1 µM from a 5 mM stock in dimethyl sulfoxide [Abcam] for 5 min); streptomycin (to non-specifically block mechano-sensitive ion channels; 50 µM from a stock concentration of 15 mM in distilled water [Sigma-Aldrich] for 5 min); allyl isothiocyanate (AITC, to activate TRPA1 channels; 10 µM from a stock concentration of 1 M in dimethyl sulfoxide [Sigma-Aldrich] for 5 min); HC-030031 (to block TRPA1 channels; 10 µM from a 50 mM stock in dimethyl sulfoxide [Abcam] for 30 min), N-acetyl-L-cysteine (NAC, to chelate ROS; 10 mM from a 0.5 M stock in distilled water [Sigma-Aldrich] for 20 min); and diphenyleneiodonium (DPI, to block ROS production; 3 µM from a 10 mM stock in dimethyl sulfoxide [Abcam] for 60 min).

### Simultaneous V_m_-Ca^2+^ Fluorescence Imaging

Whole hearts were stained with V_m_ and [Ca^2+^]_i_-sensitive fluorescent dyes (di-4-ANBDQPQ and Rhod-2-AM) and near-simultaneous V_m_-Ca^2+^ measurement was performed by dual-excitation optical mapping with a custom, camera frame-synchronised, single camera system cropped to 550 × 350 pixels recording at 520 frames/s (Figure 3a).^26–28^ Single LV cardiomyocytes were stained with V_m_ and [Ca^2+^]_i_-sensitive fluorescent dyes (di-4-ANBDQPQ and Fluo-5F-AM) and simultaneous V_m_-Ca^2+^ measurement was performed by single-excitation, dual-emission fluorescence imaging with a single camera of 128 × 128-pixels recording at 500 frames/s, using an optical splitter system for simultaneous recording on two halves of the camera sensor (Figure 5a).^29^

**Figure 3.**
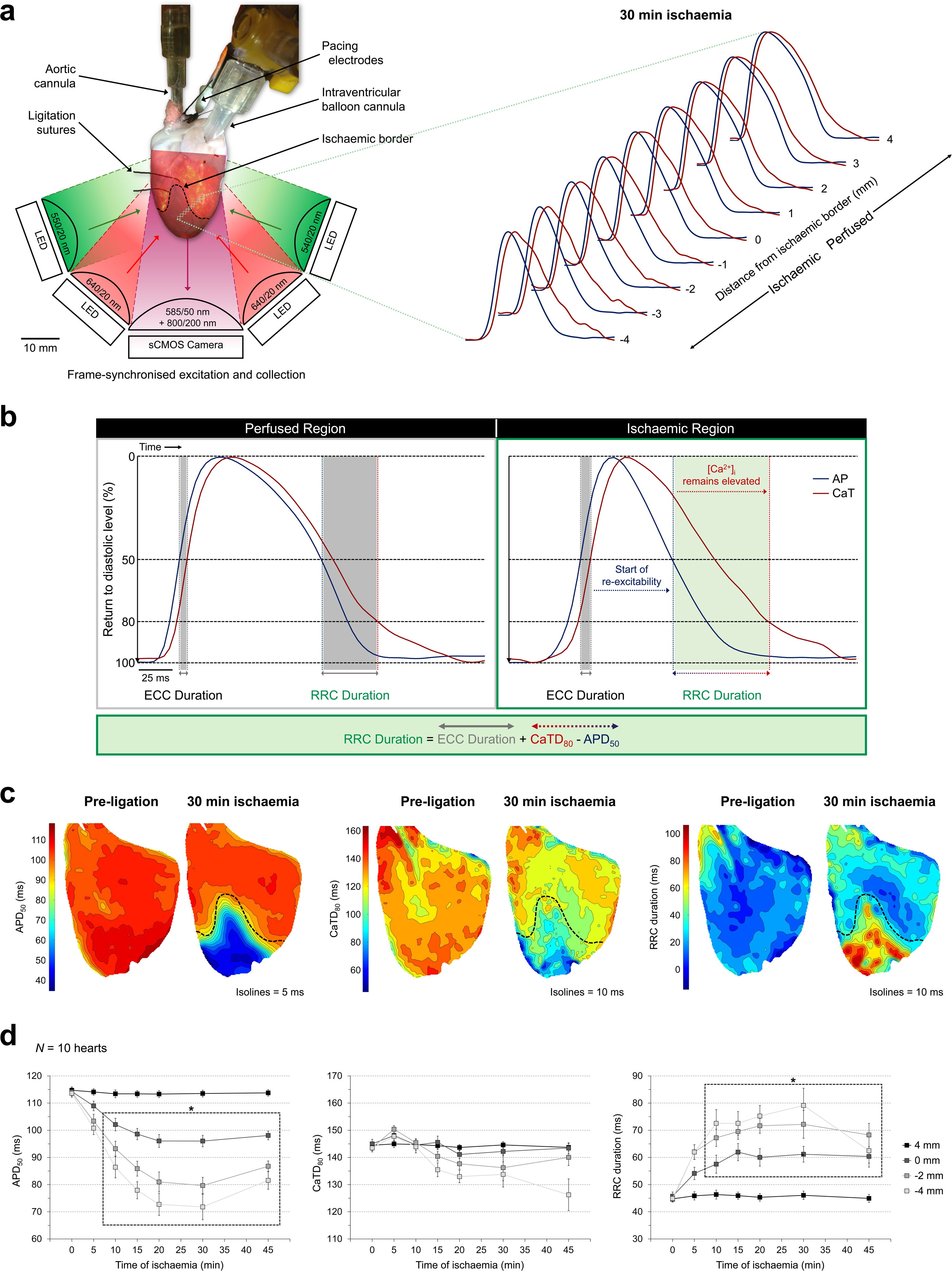
Disturbed repolarisation-relaxation coupling (RRC) during acute regional ischaemia in rabbit left ventricle (LV). **a,** Schematic illustration of the dual-excitation (using camera frame-synchronised light-emitting diodes, LED), dual-emission fluorescence imaging approach projecting the two emission wavelengths (via a 585/50 nm / 800/200 nm band-pass filter) onto a single scientific complementary metal-oxide-semiconductor (sCMOS) camera for near-simultaneous measurement of voltage and cytosolic calcium ([Ca^2+^]_i_) in rabbit whole hearts (left) and representative action potentials (AP, blue) and Ca^2+^ transients (CaT, red) along a line across the ischaemic border 30 min after coronary artery ligation (right). **b,** Representative AP and CaT traces, showing AP duration at 50% repolarisation (APD_50_, blue) and CaT duration at 80% return to diastolic levels (CaTD_80_, red), as well as the resulting RRC duration in the perfused (left, grey) and ischaemic tissue (right, green), demonstrating an ischaemia-induced prolongation of RCC duration. **c**, Maps of APD_50_ (left), CaTD_80_ (middle), and RRC duration (right) across the LV before artery ligation and 30 min after the onset of ischaemia. **d**, Measurements of APD_50_ (left) CaTD_80_ (middle) and RRC duration (right) over 45 min from artery ligation at points across the ischaemic border, with the dashed box indicating values that differed significantly from pre-ligation values. Differences compared to pre-ligation assessed by one-way ANOVA, with Dunnet *post-hoc* tests. \**p* < 0.05.

**Figure 4.**
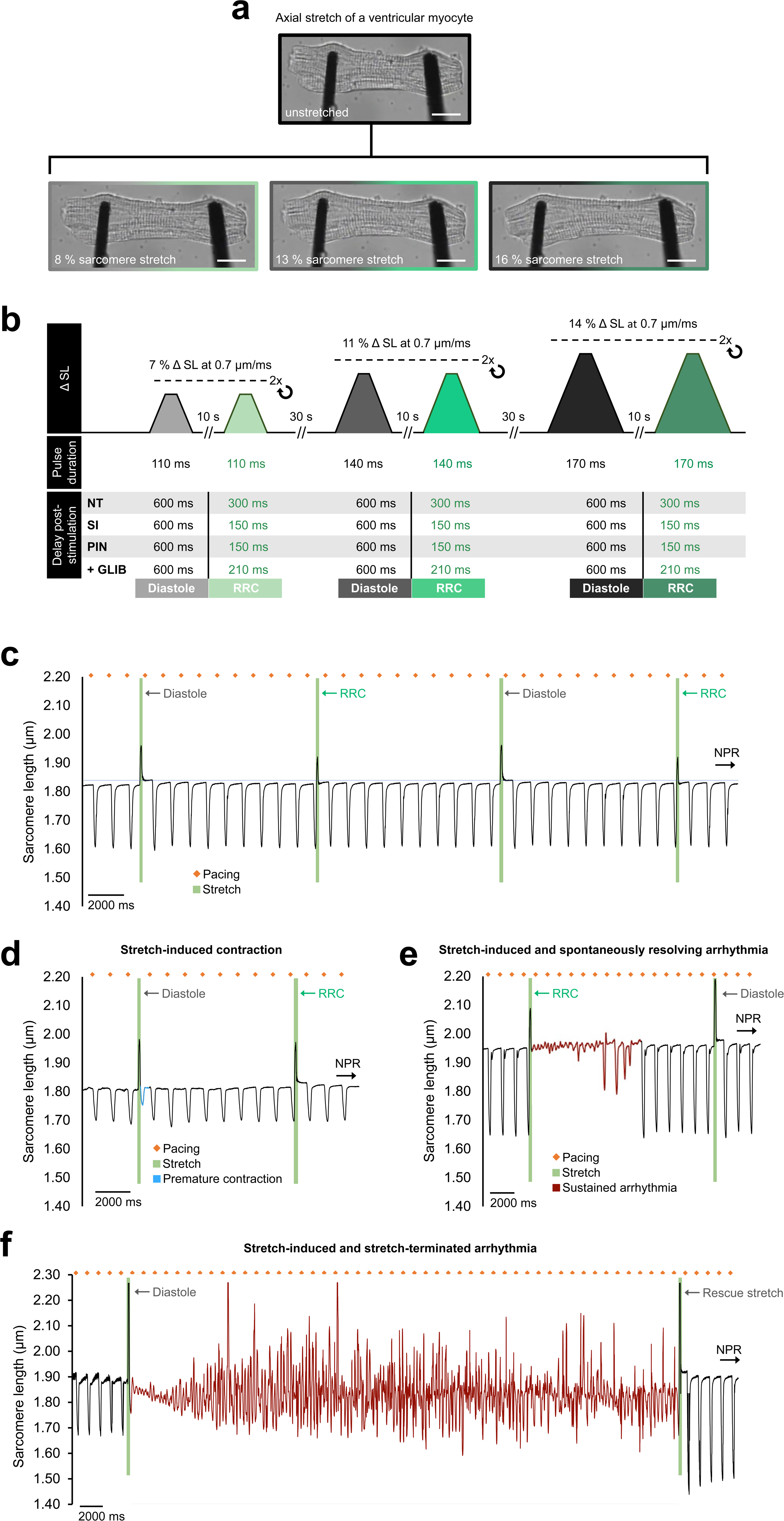
Arrhythmias elicited by transient stretch of rabbit left ventricular (LV) cardiomyocytes. **a,** Brightfield image of a rabbit LV cardiomyocyte before (top) and during (bottom) various levels of axial stretch, applied using a carbon fibre-based system. Scale bar, 10 μm. **b**, Schematic of the stretch protocol, including timing information for stretch during RRC. **c**, Representative measurement of sarcomere length in a cardiomyocyte during simulated ischaemia with 1 Hz pacing (orange dots) and stretch (green segments) applied in diastole (first and third stretch) or in late systole during repolarisation-relaxation coupling (RRC, second and fourth stretch), none of which resulted in an arrhythmia (normal paced rhythm, NPR). **d,** Stretch-induced contraction (blue curve segment) upon diastolic stretch during simulated ischaemia. **e,** Arrhythmic activity (red segment) after a stretch during RRC in simulated ischaemia conditions that spontaneously resolved. **f,** Sustained arrhythmic activity (red segment) after a diastolic stretch during pinacidil exposure, which was terminated by application of an additional stretch (rescue stretch).

**Figure 5.**
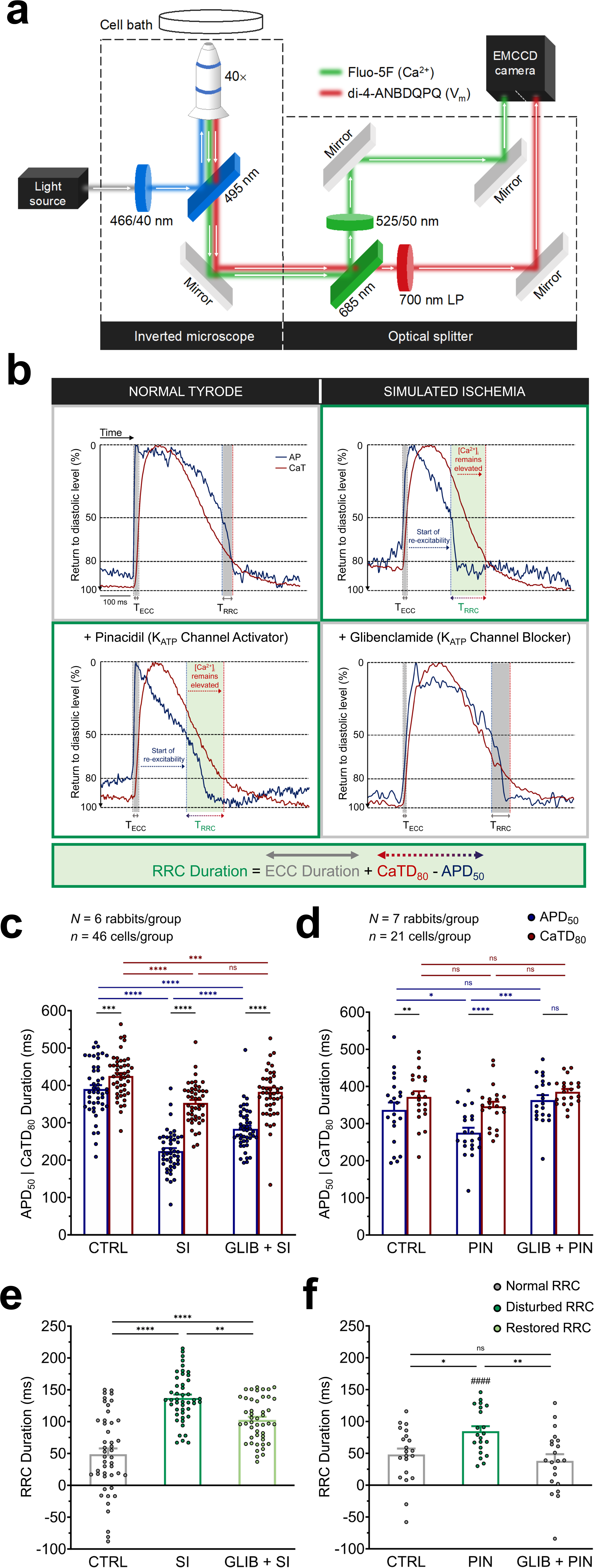
Disturbed repolarisation-relaxation coupling (RRC) in rabbit left ventricular (LV) cardiomyocytes during simulated ischaemia (SI) or ATP-sensitive potassium (K_ATP_) channel activation with pinacidil (PIN). **a**, Schematic of the single-excitation, dual-emission fluorescence imaging approach, utilising a single camera with an image splitter for simultaneous measurement of voltage and intracellular calcium ([Ca^2+^]_i_) in rabbit LV cardiomyocytes. **b,** Representative action potential (AP) and Ca^2+^ transients (CaT) showing the duration of RRC (= excitation-contraction coupling [ECC] duration + CaT duration at 80% return to diastolic levels [CaTD_80_] - AP duration at 50% repolarisation [APD_50_]) in control (CTRL, top left), during SI (top right) or PIN application (bottom left) – both of which lead to an increase in RRC duration (green shaded area) – and upon mitigation of the SI-induced RRC lengthening by glibenclamide application (GLIB, bottom right). **c,** Measurements of APD_50_ (blue) and CaTD_80_ (red) in CTRL, SI, or SI following pre-treatment with GLIB (GLIB + SI). **d,** Measurements of APD_50_ and CaTD_80_ in CTRL, or exposure to PIN or GLIB + PIN. **e,f,** RRC duration in each experimental condition. Differences between conditions assessed by one-way ANOVA, with Tukey *post-hoc* tests, and between APD_50_ and CaTD_80_ or SI and PIN by paired, two-tailed, Student’s *t*-tests. \**p* < 0.05, \*\**p* < 0.01, \*\*\**p* < 0.001, and \*\*\*\**p* < 0.0001. ^####^*p* < 0.0001 for SI *vs* PIN.

### Ratiometric [Ca^2+^]_i_ Fluorescence Imaging

Cardiomyocytes were stained with a ratiometric [Ca^2+^]_i_-sensitive fluorescent dye (Fura Red-AM), and measurement of [Ca^2+^]_i_ levels was performed by near-simultaneous collection of ratiometric signals using a custom, dual-excitation, camera frame-synchronised, single camera system.

### Tissue Slice-based Isolated Cardiomyocyte Preparation

For experiments involving patch clamp recordings of TRPA1 currents in single cells, LV cardiomyocytes were isolated from slices of LV myocardium from hearts of female New Zealand White rabbits (2.0 ± 0.2 kg, Charles River), to allow for multiple day use of rabbit LV tissue.^30^

### Sarcolemmal Ion Current Recordings

Sarcolemmal ion current was recorded by the patch-clamp technique in cell-attached configuration with protocols previously used to study cation non-selective channels.^31^

### Statistical Analysis

Values are reported as mean ± standard error of the mean. Statistical tests were performed using Prism 9 (GraphPad), with *p* < 0.05 taken to indicate statistically significant differences between groups. Applied tests are indicated in the figure captions, along with the number of replicates.

## RESULTS

### Systolic Stretch Occurs at the Ischaemic Border in Whole Hearts

Acute regional ischaemia was induced in Langendorff-perfused hearts by complete ligation of the anterior branch of the left circumflex coronary artery. After 30 min of ischaemia, in hearts in which the LV was contracting against a physiological load, this was associated with a decrease in maximum radial tissue strain (*i.e.*, reduced contraction-induced wall thickening) in the ischaemic region and at the ischaemic border (Figure 1b), indicating a local reduction in active myocardial contractile function (Figure 1c, d). There was also a significant reduction in minimum radial strain (*i.e.*, stretch-induced wall thinning) at the ischaemic border, indicating passive stretch. This occurred during LV contraction, rather than during relaxation-induced ventricular filling (Figure 1c, d).

### Mechanical Activity and Free Cytosolic Ca^2+^ are Critical for Arrhythmogenesis During Acute Ischaemia in Whole Hearts

Following coronary ligation in hearts with the LV contracting against a physiological load, ectopic excitation and/or self-terminating arrhythmic activity (Figure 2a) occurred in a bimodal fashion over 60 min of ischaemia (as shown previously in patients and preclinical models),^11,32^ with the first period of arrhythmogenesis between 0-20 min (phase 1a) and a second between 20-45 min (phase 1b, Figure 2b). To assess the importance of mechanical activity for the observed arrhythmias, hearts were either mechanically unloaded (by emptying the intraventricular balloon) or made non-contracting but still electrically active (by application of the excitation contraction uncoupler blebbistatin). In both cases, reducing mechanical activity was accompanied by a diminished arrhythmia incidence (Figures 2c, d). Next, as: (i) during ischaemia altered Ca^2+^ handling results in elevated [Ca^2+^]_i_,^33^ which can be arrhythmogenic by driving forward-mode sodium-Ca^2+^-exchanger (NCX) activity and by shifting cardiomyocyte V_m_ closer to the threshold for excitation,^34^ and (ii) a stretch-induced influx of sodium and/or Ca^2+^ will directly cause V_m_ depolarisation and potentially further contribute to NCX-mediated inward currents,^6^ the importance of [Ca^2+^]_i_ for arrhythmogenesis was investigated by buffering with BAPTA in contracting hearts with a physiologically loaded LV. As in the case of reduced mechanical activity, [Ca^2+^]_i_ buffering was associated with a pronounced reduction in arrythmia incidence during the 60 min of regional ischaemia (Figures 2e). Since [Ca^2+^]_i_ has been shown to rise during acute ischaemia by increased leak from the sarcoplasmic reticulum *via* ryanodine receptors,^35^ and as stretch is known to also cause ryanodine receptor-mediated Ca^2+^ leak^7^ (with this mechano-sensitive response being enhanced in ischaemia^23^), we explored whether stretch-induced sarcoplasmic reticulum Ca^2+^ leak may also contribute to observed arrhythmias by stabilising ryanodine receptors with dantrolene. As during exposure to BAPTA, dantrolene reduced arrythmia incidence (Figure 2f). When arrhythmia incidence during each intervention was assessed for the two distinct ischaemia periods (phase 1a and 1b) and compared to the incidence in contracting hearts with a physiologically loaded LV, a significant decrease was found in hearts with a mechanically unloaded LV and in non-contracting hearts during both phases, while [Ca^2+^]_i_ buffering and ryanodine receptor stabilisation gave rise to a significant reduction only in phase 1b (Figure 2g, Figure S2).

While the above results suggest that mechanical activity and high [Ca^2+^]_i_ levels are critical for arrhythmogenesis during acute ischaemia in the whole heart, the reduced incidence of arrhythmias with application of BAPTA or dantrolene may have been a secondary effect of Ca^2+^ buffering on contractile function. This possibility was assessed by measuring the effect of each drug on the maximum rate of LV pressure development and on LV developed pressure during contraction. BAPTA application had no significant effect on either measure, while dantrolene reduced both (Figure S3), suggesting that the reduction in arrhythmogenesis with dantrolene may have been due, at least in part, to reduced mechanical activity.

### RRC is Disturbed in Ischaemic Tissue in Whole Hearts

As [Ca^2+^]_i_ dynamics appeared to be involved in arrhythmogenesis during phase 1b of acute ischaemia in contracting hearts with a physiologically loaded LV (Figure 2e), changes in [Ca^2+^]_i_ dynamics in relation to V_m_ were investigated by simultaneous V_m_-Ca^2+^ optical mapping (Figure 3a). To characterise ischaemia-induced changes in RRC, a time window was defined to capture changes in post-activation recovery of [Ca^2+^]_i_ and V_m_. This was calculated as the sum of excitation-contraction coupling duration (the time from start of the AP to the start of the CaT) plus CaTD at 80% recovery (CaTD_80_), minus APD at 50% repolarisation (APD_50_), and allows one to track the period during which cardiomyocytes begin to become re-excitable while [Ca^2+^]_i_ remains elevated (Figure 3b). After 10 min of acute ischaemia, APD_50_ in the ischaemic region and at the ischaemic border was reduced, compared to pre-ligation values, while CaTD_80_ was not significantly altered, resulting in an increase in RRC duration (there were no significant changes in the perfused tissue; Figure 3c, d).

### Transient Stretch Triggers Mechano-Arrhythmogenesis in Cardiomyocytes

To investigate whether cardiomyocyte stretch might account for the arrhythmias seen during acute regional ischaemia in the whole heart, carbon fibres were attached to both ends of single LV cardiomyocytes and axial stretch was applied under piezo-electric actuator control (Figure 4a). Systolic (during RRC) and diastolic stretch (Figure 4b) resulted in a variety of disturbances in cardiomyocyte mechanical activity (revealed by tracking sarcomere length; Figure 4c), including stretch-induced contractions (sometimes followed by a second contraction out of sync with the continuing electrical stimulation; Figure 4d and Video S3) and self-sustained rhythm disturbances (3 or more successive contractions immediately after stretch) which either resolved spontaneously (Figure 4e) or were terminated by application of an additional stretch (Figure 4f, Video S4).

Since previous work in the whole heart has shown that the magnitude of tissue deformation is a key determinant of mechano-arrhythmogenesis,^36^ we assessed whether incidence of stretch-induced beating irregularities in cardiomyocytes is also dependent on stretch amplitude. Increasing piezo-electric actuator displacement step amplitudes from 20 to 30 or 40 μm resulted in an increase in the percent sarcomere stretch (20 μm: 7.0 ± 0.5%; 30 μm: 11.1 ± 0.7%; 40 μm: 14.4 ± 0.9%) and the maxima of sarcomere length (20 μm: 1.89 ± 0.02 μm; 30 μm: 1.96 ± 0.02 μm; 40 μm: 2.02 ± 0.03 μm) and applied stress (20 μm: 2.95 ± 0.04 mN/mm^2^; 30 μm: 3.88 ± 0.05 mN/mm^2^; 40 μm: 5.00 ± 0.04 mN/mm^2^) during stretch pulses in all cardiomyocytes (Figures 4a and S4). The increase in applied stretch corresponded with an increased incidence of beating irregularities (Figure S5), illustrating that mechano-arrhythmogenesis in cardiomyocytes, as in the whole heart, is affected by stretch magnitude.

### K_ATP_ Channel Activation Disturbs RRC in Cardiomyocytes

To mimic ischaemic changes after 30 min of ischaemia in the whole heart, LV cardiomyocytes were exposed to: (i) SI solution, which combined metabolic inhibition (block of oxidative phosphorylation and anaerobic glycolysis) with acidosis and hyperkalemia,^11,23,24^ or (ii) solution containing PIN, which is a well-established agonist of sulfonylurea receptor 2A / K_ir_6.2 (the hetero-octamer that forms K_ATP_ channels in cardiac and skeletal muscle^37^), to assess the isolated effects of ischaemia-induced K_ATP_ channel opening. The dyes di-4-ANBDQPQ (for V_m_) and Fluo-5F-AM (whose relatively high K_d_ value [∼2.3 μM] limits its buffering of diastolic [Ca^2+^]_i_, and thus helps to avoid potential artefactual lengthening of the CaTD) were used to simultaneously measure V_m_ and [Ca^2+^]_i_ transients, and to assess RRC duration in electrically-paced rabbit LV cardiomyocytes under carbon fibre control (Figure 5a).

Under physiological conditions with normal RRC (in CTRL solution), CaTD_80_ was significantly longer than APD_50_ (individually true in 80% of cases; Figure 5b-d). During exposure to SI or PIN, APD_50_ decreased (Figure 5b-d), with a greater decrease occurring during SI (suggesting a contribution of factors beyond K_ATP_ channel activation to APD shortening^11^). CaTD_80_ also decreased during SI (as can be expected with metabolic inhibition and acidosis^33^), but to a lesser extent than APD_50_, whereas with PIN there was no significant change in CaTD_80_. As a result, RRC was prolonged during both SI and PIN (Figure 5b, e-f), but to a greater extent during SI, due to the greater relative decrease in APD_50_ than CaTD_80_ compared to PIN. These changes in APD_50_ and CaTD_80_ mimic the behaviour seen in the whole heart during acute regional ischaemia (Figure 3). In cardiomyocytes pre-treated with glibenclamide to block K_ATP_ channels, the reduction in APD_50_ was attenuated (during SI) or prevented (during PIN), with the greater effect during PIN again suggesting K_ATP_-independent contributions to APD shortening during SI. In contrast, there was no significant effect of glibenclamide on CaTD_80_ (Figure 5b-d), such that RRC duration decreased compared to either SI or PIN alone (Figure 5b, e, f).

### Disturbed RRC Permits Systolic Mechano-arrhythmogenesis in Cardiomyocytes

To determine whether an increased RRC duration may enable systolic mechano-arrhythmogenesis in LV cardiomyocytes, stretch protocols were repeated with cells exposed to SI or to PIN. In both cases, there was an increase in the incidence of mechano-arrhythmogenesis with stretch in late systole compared to CTRL (Figure 6a) and, as in CTRL, an increase in stretch magnitude (quantified as percent sarcomere stretch, maximum sarcomere length during stretch, and maximum applied stress; Figure S4) correlated with the incidence of beating irregularities (Figure S5). Importantly, there was no difference in the percent sarcomere stretch, maximum sarcomere length during stretch, or maximum applied stress across groups (CTRL, SI, or PIN; Figure S4), suggesting that there were no significant changes in passive mechanical properties of cardiomyocytes, or subsequent changes in characteristics of the mechanical stimulus cells experienced, that would account for the increased incidence of mechano-arrhythmogenesis during SI or PIN exposure.

**Figure 6.**
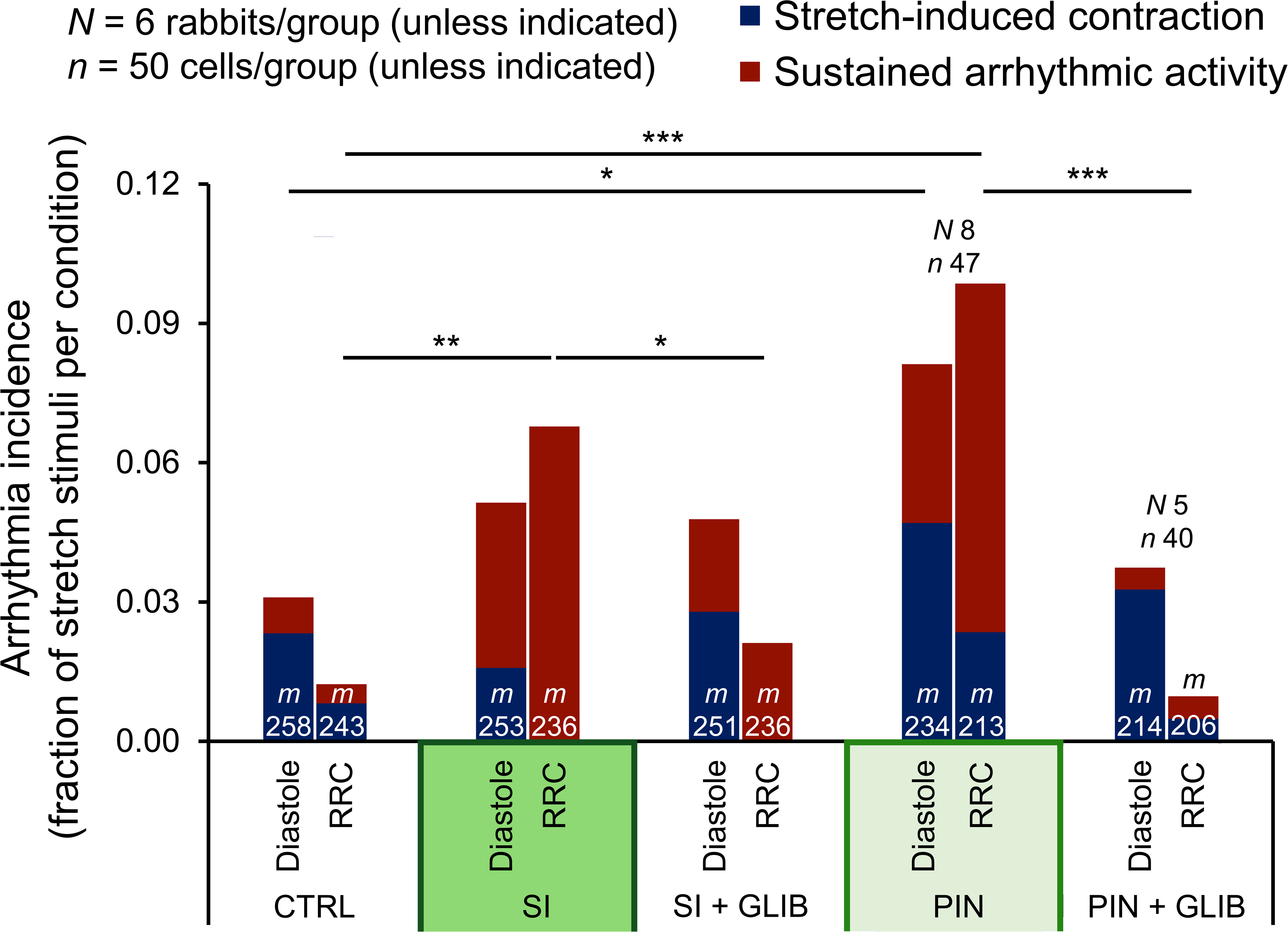
Role of disturbed repolarisation-relaxation coupling (RRC) in mechano-arrhythmogenesis. Incidence of stretch-induced contractions (blue) and self-sustained arrhythmias (red) with stretch of rabbit left ventricular cardiomyocytes, applied during diastole or RRC, in control conditions (CTRL), or during exposure to simulated ischaemia (SI), glibenclamide (GLIB) + SI, pinacidil (PIN), or GLIB + PIN. Differences in arrhythmia incidence assessed using chi-square contingency tables and Fisher’s exact test. \**p* < 0.05, \*\**p* < 0.01, and \*\*\**p* < 0.001 between groups. *m* = stretch stimuli applied.

To test whether preventing the SI- or PIN-induced prolongation of RRC may affect the increased probability of systolic mechano-arrhythmogenesis, cardiomyocytes were pre-treated with glibenclamide. Glibenclamide (which reduces [SI] or prevents [PIN] the increase in RRC duration; Figure 5b, e, f) prevented the increase in systolic mechano-arrhythmogenesis in both groups (Figure 6), suggesting that the presence of physiological RRC durations protects cardiomyocytes from arrhythmogenesis, while disturbed RCC raises susceptibility to mechano-arrhythmogenesis during late systole. Of note, glibenclamide pre-treatment had no significant effects on diastolic mechano-arrhythmogenesis in cardiomyocytes exposed to PIN (Figure 6).

SI had no significant effects on the incidence of mechano-arrhythmogenesis with stretch in diastole compared to CTRL, indicating that ischaemic changes in RRC do not affect diastolic mechano-arrhythmogenesis. In contrast, there was an increased incidence of beating irregularities with diastolic stretch in PIN-treated cardiomyocytes (Figure 6a). This may be related to secondary effects of PIN, which were explored in additional experiments, described in the following section.

To ascertain whether stretch-induced cell damage could account for the observed increases in mechano-arrhythmogenesis, cardiomyocyte contractile function was assessed before and after stretch of each magnitude. In cardiomyocytes exposed to CTRL, SI, or PIN, diastolic sarcomere length and the maximum rate and percent of sarcomere shortening were not significantly different before and after each round of stretch applications, suggesting that cardiomyocyte damage or carbon fibre slippage during stretch are unlikely to have occurred (Figure S6). Recorded parameters were also not significantly different before and after a return to steady state following periods of sustained rhythm disturbances, irrespective of arrhythmia duration (Figure S7), indicating that cellular damage is also unlikely to underlie the more severe forms of mechano-arrhythmogenesis.

### [Ca^2+^]_i_ Plays a Role in Mechano-Arrhythmogenesis in Cardiomyocytes

Given that in hearts with a contracting, physiologically loaded LV, [Ca^2+^]_i_ is critical for arrhythmogenesis during acute regional ischaemia (Figure 2e), we sought to assess whether [Ca^2+^]_i_ also plays a role in cardiomyocyte-level mechano-arrhythmogenesis. As in the whole heart, [Ca^2+^]_i_ was buffered with BAPTA, which has previously been shown in ventricular cardiomyocytes to counter stretch-induced changes in cardiac electrophysiology driven by increases in [Ca^2+^]_i_.^38^ In cardiomyocytes with an SI- or PIN-induced prolongation of RRC, BAPTA reduced the incidence of systolic mechano-arrhythmogenesis (Figure 6a; in CTRL, application of BAPTA had no significant effect; Figure S8). Ca^2+^ buffering also prevented the increase in diastolic mechano-arrhythmogenesis (Figure 7a) that occurs in PIN-treated cells (Figure 6a). Against expectations, BAPTA increased the incidence of diastolic mechano-arrhythmogenesis in cardiomyocytes exposed to SI (Figure 7a), in particular of single stretch-induced beats; this may be explained by a Ca^2+^ buffering-induced increase in the trans-sarcolemmal driving force for Ca^2+^ entry during stretch, specifically in ischaemic conditions.

**Figure 7.**
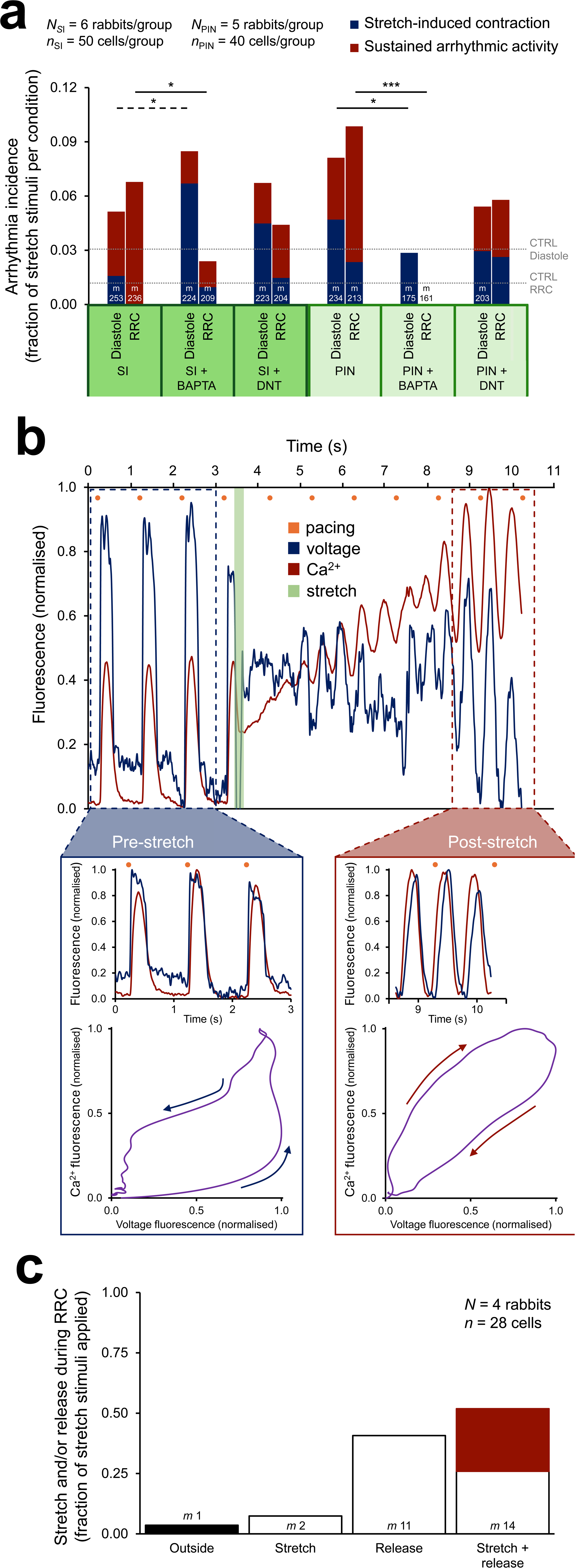
Contribution of cytosolic calcium concentration ([Ca^2+^]_i_) to mechano-arrhythmogenesis. **a,** Incidence of stretch-induced contractions (blue) and sustained arrhythmias (red) with stretch of rabbit left ventricular (LV) cardiomyocytes, applied either during diastole or during repolarisation-relaxation coupling (RRC), upon exposure to simulated ischaemia (SI) or pinacidil (PIN), as well as in combination with BAPTA (to buffer [Ca^2+^]_i_) or dantrolene (DNT, to stabilise ryanodine receptors in their closed state). Dashed lines show arrhythmia incidence for stretch during diastole or RRC in control (CTRL) conditions. **b,** Representative voltage (blue) and Ca^2+^ (red) signals simultaneously recorded by fluorescence imaging in a cardiomyocyte exposed to PIN, showing a sustained arrhythmia, induced by stretch during the period of prolonged RRC (green), leading initially to depolarisation and an associated increase in [Ca^2+^]_i_ above normal systolic levels, followed by spontaneous oscillations in [Ca^2+^]_i_ (which subsequently resolved, followed by normal paced beats; not shown). **Blue inset,** Scaled voltage and Ca^2+^ signals (top) and phase plot of the first beat in the insert (bottom), showing that prior to stretch, changes in voltage precede changes in [Ca^2+^]_i_. **Red inset,** Scaled voltage and Ca^2+^ signals (top) and phase plot of the first beat in the insert (bottom), showing [Ca^2+^]_i_ oscillations preceding changes in voltage during the stretch-induced sustained arrhythmia, suggesting aberrant Ca^2+^ handling as the driving mechanism. **c,** Fraction of stretch stimuli applied for which stretch and/or release occurred during RRC in a subset of cells exposed to SI. Stimuli that resulted in arrhythmias are shown in red. Differences in arrhythmia incidence assessed using chi-square contingency tables and Fisher’s exact test. \**p* < 0.05 and \*\*\**p* < 0.001 between groups (dashed line indicates an increase). *m* = stretch stimuli applied.

Ryanodine receptor stabilisation with dantrolene decreased the incidence of arrythmias during acute regional ischemia in the whole heart (Figure 2f), but it was unclear whether this may partly have been due to effects on mechanical activity (Figure S2). We therefore exposed LV cardiomyocytes to dantrolene (as in the whole heart) before stretch (which has been shown not to affect Ca^2+^-induced Ca^2+^ release or contractile function in single cardiomyocytes^39^). Dantrolene had no significant effect on the incidence of mechano-arrhythmogenesis in cardiomyocytes exposed to SI or PIN (Figure 7a; in CTRL there was also no significant effect, Figure S8), suggesting that effects on mechanical activity may indeed have been partly responsible for the reduced arrhythmia incidence in the whole heart.

To assess whether the reduction in mechano-arrhythmogenesis with BAPTA was a result of secondary effects on cardiomyocyte mechanics, contractile function was assessed. Treatment of CTRL cells with BAPTA (as well as dantrolene) had no effect on diastolic sarcomere length or maximum rate of systolic sarcomere shortening; there was a moderate decrease in the percent systolic sarcomere shortening with BAPTA, in keeping with a buffering effect on peak systolic [Ca^2+^]_i_ (Figure S9).

The above findings confirm that [Ca^2+^]_i_ is important for mechano-arrhythmogenesis in cardiomyocytes with disturbed RRC. The most definitive evidence for a direct role of [Ca^2+^]_i_ in mechano-arrhythmogenesis came from dual V_m_-Ca^2+^ fluorescence imaging of cardiomyocytes during sustained rhythm disturbances. This revealed progressive increases in [Ca^2+^]_i_ after stretch application, which were followed by Ca^2+^ oscillations that preceded changes in V_m_, indicating that [Ca^2+^]_i_ fluctuations can drive stretch-triggered arrhythmias (Figure 7b, Video S5).

### Systolic Mechano-Arrhythmogenesis Depends on Stretch and Release in Cardiomyocytes

While systolic arrhythmias can be elicited with stretch occurring during RRC, termination of stretch (‘release’) may also be arrhythmogenic.^8^ Stretch of cardiac tissue increases the affinity of myofilaments for Ca^2+^, such that stretch during systole, when [Ca^2+^]_i_ is high, results in increased Ca^2+^ binding. Upon release, rapid dissociation of myofilament-bound Ca^2+^ may produce a surge in [Ca^2+^]_i_, which can trigger Ca^2+^ release, Ca^2+^ waves, or NCX-mediated membrane depolarisation.^8^

In a subset of cardiomyocytes exposed to SI, fluorescence-based measurements of APD and CaTD during late-systolic stretch was performed to assess the phases of the stretch pulse (*i.e.,* stretch, release, or stretch-and-release) that occurred during RRC, and relate this to mechano-arrhythmogenesis. The analysis confirmed that in 96% of cases, at least part of the stretch pulse was timed to occur during the cardiomyocyte-specific RRC period. However, arrhythmias occurred only if both stretch *and* release occurred during RRC; this timing was present in just over half of the cells tested. In these cardiomyocytes, mechano-arrhythmogenesis was observed in 50% of cases (*i.e.*, in about 25% of RRC-targeted stretch applications; Figure 7c). These results suggest that the combination of stretch-and-release may be particularly prone to causing late-systolic mechano-arrhythmogenesis, and confirms the successful targeting of mechanical stimulation to this period in the experiments presented above.

### Activated TRPA1 Channels Drive Mechano-Arrhythmogenesis in Cardiomyocytes

While it was anticipated that prolonged RRC in cardiomyocytes during exposure to SI or to PIN would raise the incidence of systolic mechano-arrhythmogenesis, the increased incidence of diastolic arrhythmogenesis with PIN (Figure 6a) is surprising. PIN is generally considered a specific agonist of K_ATP_ channels, resulting in a repolarising efflux of potassium from cardiomyocytes. However, in HEK293 cells, PIN has been shown to increase the activity of TRPA1 channels, resulting in a depolarising influx of Ca^2+^.^40^ This is potentially important in the context of mechano-arrhythmogenesis, as TRPA1 channels: (i) are inherently mechano-sensitive^41^ and contribute to mechanically-evoked electrical responses in sensory neurons,^42–45^ astrocytes,^46^ and vertebrate hair cells;^47^ (ii) are functionally expressed in *Drosophila* hearts^48^ and murine ventricular cardiomyocytes;^49^ and (iii) are important for cardiovascular function and disease progression.^50^

To explore whether TRPA1 channels could contribute to the increase in systolic and diastolic mechano-arrhythmogenesis upon PIN treatment in rabbit LV cardiomyocytes, ion channel activity was measured in cell-attached patches during application of the TRPA1 channel-specific agonist, AITC.^51^ AITC exposure caused an increase in total inward current (with a ∼1-3 min delay, similar to the time-lag in response reported in other studies^52–53^), while there was no change in time-matched CTRL cells (Figure 8a, b). To explore whether activated TRPA1 channels can contribute to mechano-arrhythmogenesis, cardiomyocytes were exposed to AITC (which has been shown to also enhance the response of TRPA1 channels to mechanical stimulation^44^) and subjected to diastolic stretch. Cardiomyocytes exposed to AITC showed an increased incidence of diastolic mechano-arrhythmogenesis compared to CTRL, which was prevented by application of BAPTA, as well as by application of the specific TRPA1 channel antagonist HC-030031^54^ (which has been shown to inhibit TRPA1 current and mechanically-induced excitation in sensory neurons;^42–45^ Figure 8c).

**Figure 8.**
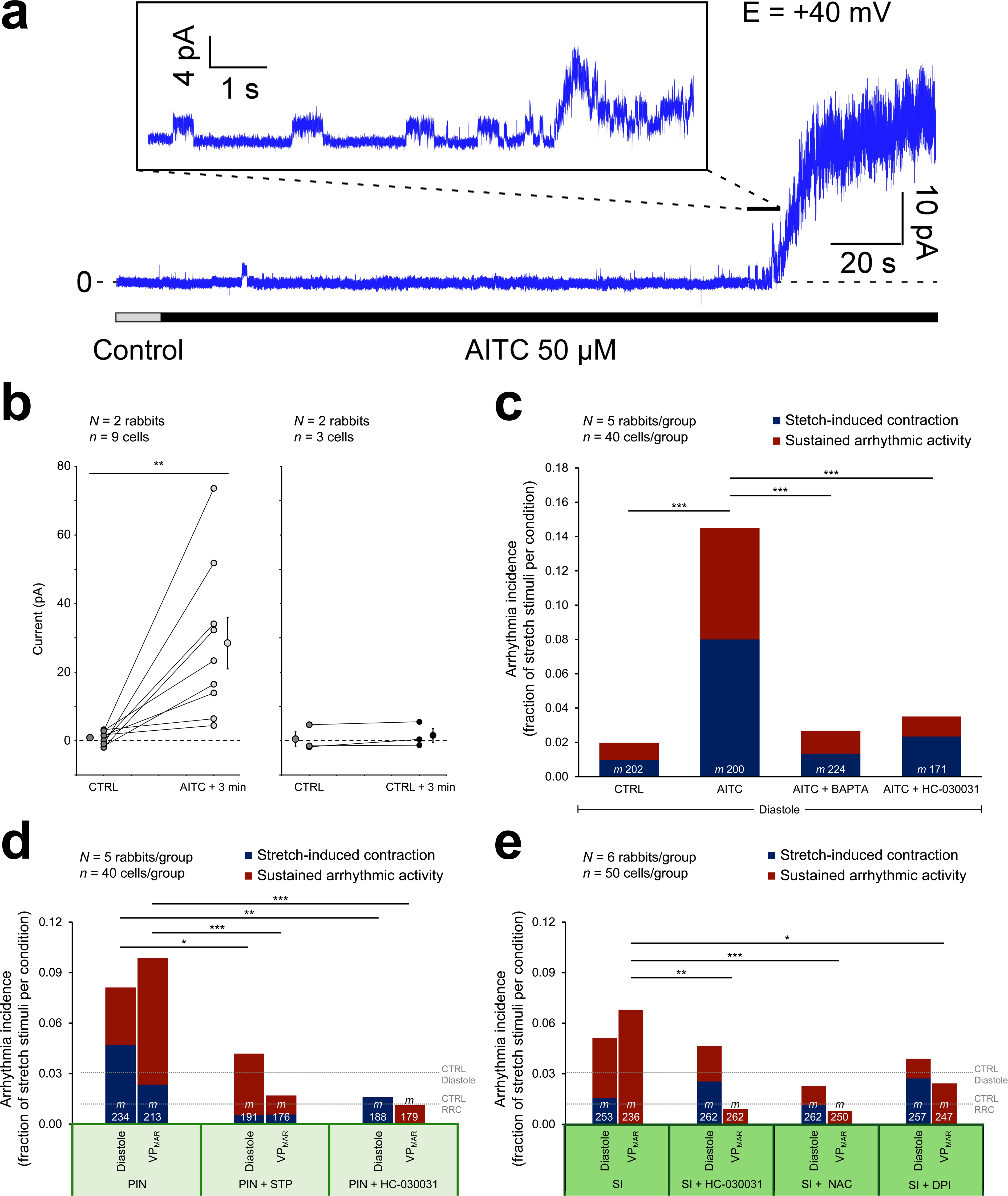
Contribution of transient receptor potential kinase ankyrin 1 (TRPA1) channels to mechano-arrhythmogenesis. **a,** Representative current recording by cell-attached patch clamp (holding potential +40 mV) of rabbit left ventricular (LV) cardiomyocytes in control (CTRL) and after switching to allyl isothiocyanate (AITC)-containing solution (50 μM; to activate TRPA1 channels). **Inset,** Detail of channel activation with visible single channel events. **b**, Quantification of AITC-induced current changes (left; 3 min total application including a 15 s lag time for AITC to reach the cardiomyocyte) and time-matched, same-batch cardiomyocytes in CTRL conditions (right; 3 min). **c,** Incidence of stretch-induced contractions (blue) and self-sustained arrhythmias (red) after stretch of cardiomyocytes during diastole in CTRL and during exposure to AITC, AITC + BAPTA (to buffer cytosolic calcium concentration, [Ca^2+^]_i_), or AITC + HC-030031 (to block TRPA1 channels). **d**, Incidence of mechano-arrhythmogenesis upon stretch of cardiomyocytes during diastole or RRC during exposure to pinacidil (PIN), PIN + streptomycin (STP, a non-specific blocker of mechano-sensitive ion channels), or PIN + HC-030031. **e**, Incidence of mechano-arrhythmogenesis during exposure to simulated ischaemia (SI), or to SI with additional exposure to HC-030031, N-acetyl-L-cysteine (NAC, to chelate reactive oxygen species, ROS), or diphenyleneiodonium (DPI, to block ROS production). Dashed lines in **d** and **e** show arrhythmia incidence for stretch during diastole or RRC in control (CTRL) conditions. Current differences assessed with two-tailed, paired, Student’s *t*-test. \*\**p* < 0.01. Differences in arrhythmia incidence assessed using chi-square contingency tables and Fisher’s exact test. \**p* < 0.05, \*\**p* < 0.01, and \*\*\**p* < 0.001 between groups. *m* = stretch stimuli applied.

To assess whether TRPA1 channels contribute to the increase in systolic and diastolic mechano-arrhythmogenesis with PIN, LV cardiomyocytes were subjected to PIN or to PIN with either streptomycin (a non-specific antagonist of cation non-selective mechano-sensitive ion channels^55^) or the TRPA1-specific antagonist HC-030031 and subjected to stretch during diastole or RRC. Streptomycin and HC-030031 both reduced the incidence of mechano-arrhythmogenesis upon diastolic and late systolic stretch in PIN-treated cardiomyocytes (Figure 8d; arrhythmia incidence was not significantly different in CTRL and CTRL + HC-030031 conditions, Figure S8; neither drug had any other significant effect on functional assays, Figure S9), supporting the suggestion that TRPA1 channels may contribute to the increased incidence of diastolic and systolic mechano-arrhythmogenesis caused by PIN.

As TRPA1 channels have been shown to be activated in ischaemic conditions,^56,57^ their contribution to the increase in systolic mechano-arrhythmogenesis seen during SI was also investigated. It was found that blocking TRPA1 channels with HC-030031 reduced systolic mechano-arrhythmogenesis during SI in isolated cardiomyocytes (Figure 8e).

To assess whether the reduction in mechano-arrhythmogenesis with streptomycin or HC-030031 and the increase with AITC might have been the result of secondary effects on cardiomyocyte mechanics, contractile function during drug treatment was assessed. In all cases, there were no significant effects on diastolic sarcomere length and maximum rate or percent of systolic sarcomere shortening (Figure S9).

### ROS Mediates Mechano-Arrhythmogenesis in Cardiomyocytes during Ischaemia

TRPA1 channels are known to be activated by an increase in [Ca^2+^]_i_^58^ or ROS,^59^ both of which are increased during ischaemia.^11^ Experiments using BAPTA to buffer [Ca^2+^]_i_ suggested a contribution of [Ca^2+^]_i_ to systolic mechano-arrhythmogenesis in SI, which may in part involve activation of TRPA1. To determine whether ROS also contributes to the ischaemia-induced increase in systolic mechano-arrhythmogenesis, cardiomyocytes were incubated with NAC (to chelate intracellular ROS) or DPI (to inhibit NOX2-mediated mechano-sensitive ROS production, which is enhanced in ischaemia^23^) prior to SI. The incidence of systolic mechano-arrhythmogenesis was decreased by both NAC and DPI pre-treatment (Figure 8e), suggesting a contribution of ROS to systolic mechano-arrhythmogenesis during SI. The contribution of ROS to mechano-arrhythmogenesis may act *via* a ROS-mediated increase in TRPA1 activity (enhancing its mechano-sensitive depolarising current and contributing to increased [Ca^2+^]_i_^60^), or through a TRPA1-independent increase in [Ca^2+^]_i_.^61^

### TRPA1 and [Ca^2+^]_i_ each can Drive Mechano-Arrhythmogenesis in Cardiomyocytes

Combined, the above results suggest that increased mechano-sensitive TRPA1 channel activity and increased [Ca^2+^]_i_ contribute to a higher incidence of mechano-arrhythmogenesis during SI and PIN-induced K_ATP_ activation. However, as it had been shown that TRPA1 itself can lead to increased [Ca^2+^]_i_,^60^ it remained unclear whether the two mechanisms may independently contribute to mechano-arrhythmogenesis. [Ca^2+^]_i_ was measured by ratiometric fluorescence imaging, which showed that diastolic [Ca^2+^]_i_ increased during exposure to SI, PIN, or AITC. This increase was reduced by TRPA1 channel block with HC-030031 during exposure to AITC only (Figure S10), suggesting that TRPA1 channels are primary drivers of an increase in [Ca^2+^]_i_ during activation with AITC, while with SI or PIN the increase is driven by other mechanisms (Figure S10; of note: in SI, chelating ROS with NAC also did not prevent the increase in diastolic [Ca^2+^]_i_). As blocking TRPA1 channels with HC-030031 (Figure 8c-e) and buffering of [Ca^2+^]_i_ with BAPTA (Figure 7a, 8c) decreased mechano-arrhythmogenesis during exposure to SI, PIN, and AITC, it would appear that increased TRPA1 activity and increased [Ca^2+^]_i_ content can independently contribute to increased mechano-arrhythmogenesis (unfortunately, the effects of BAPTA on [Ca^2+^]_i_ could not be measured, as the combined buffering of Ca^2+^ by BAPTA and the reporter Fura Red decreases cardiomyocyte contractile function).

## DISCUSSION

Taken together, the results from this study demonstrate that myocardial ischaemia disturbs RRC, and that this results in an increase in the incidence of systolic mechano-arrhythmogenesis in the whole heart and in single LV cardiomyocytes, while normal RRC or prevention of RRC prolongation (by K_ATP_ channel antagonism) is protective. Key molecular mechanisms driving the ischaemia-induced increase in systolic mechano-arrhythmogenesis are activation of TRPA1 channels and an increase in [Ca^2+^]_i_, both of which can be exacerbated by an increase in ROS.

Key findings suggest contributions of disturbed RRC, K_ATP_ and TRPA1 channels, [Ca^2+^]_i_, and ROS to the triggering (*via* a depolarising trans-sarcolemmal current) and sustenance (*via* the generation of a [Ca^2+^]_i_-mediated substrate) of mechano-arrhythmogenesis in ischaemic LV cardiomyocytes. Triggered activity during ischaemia may be driven by Ca^2+^ influx through mechano-sensitive TRPA1 channels.^41^ In ischaemia, TRPA1 channels may be persistently activated by increased [Ca^2+^]_i_^58^ and ROS.^59^ Stretch is known to increase the trans-sarcolemmal influx of cations through activated TRPA1 (which preferentially pass Ca^2+^ in HEK293 cells^62^), directly leading to V_m_ depolarisation.^42–47^ At the same time, activation of TRPA1 channels will increase [Ca^2+^]_i_, leading to an increase in the Ca^2+^ bound by the myofilaments during stretch and contributing to a greater surge of [Ca^2+^]_i_ with its dissociation during release. Increased [Ca^2+^]_i_ may cause further V_m_ depolarisation *via* electrogenic forward-mode NCX activity.^34^ If the resulting V_m_ depolarisation from these various sources is sufficiently large, it will trigger cardiomyocyte excitation and contraction. This is particularly arrhythmogenic when both stretch and release occur during RRC – a period during which [Ca^2+^]_i_ remains high in progressively re-excitable cardiomyocytes – which is facilitated in ischaemia by RRC prolongation with K_ATP_ channel activation. The resulting triggered activity may devolve into sustained arrhythmias if [Ca^2+^]_i_ stays sufficiently elevated, creating an arrhythmogenic substrate.^34^ That the combination of stretch-and-release during RRC appears to be particularly arrhythmogenic may be one reason why the overall incidence of mechano-arrhythmogenesis is not higher; only in ∼50% of ischaemic cells stretch-and-release was fully contained within the RRC period, and in only ∼50% of those cases arrhythmias were induced. The apparent importance of a stretch-release related surge of [Ca^2+^]_i_ is in keeping with prior reports^8^ and may be further explored in future experiments with pharmacological modification of myofilament Ca^2+^ sensitivity.

We further found that exposure of LV cardiomyocytes to AITC causes direct activation of TRPA1 channels and subsequent diastolic [Ca^2+^]_i_ loading, while PIN either directly activates TRPA1 channels or does so indirectly *via* an increase in [Ca^2+^]_i_. During SI, TRPA1 channels were most likely activated by elevated [Ca^2+^]_i_ or ROS. However, increased TRPA1 activity in ischaemia may also involve a change in Ca^2+^-mediated channel inhibition, or a shift in voltage-dependent gating to physiologically relevant values (half-maximal TRPA1 channel activation normally occurs between +90 and +170 mV, but is shifted as low as -36 mV by agonists such as [Ca^2+^]_i_ or ROS).^63^ Accordingly, reducing RRC duration, blocking TRPA1 channels, and limiting diastolic [Ca^2+^]_i_ or ROS reduced the incidence of systolic mechano-arrhythmogenesis. Additionally, K_ATP_ channels have been shown to be mechano-sensitive.^64^ While quiescent under physiological conditions, if activated through a reduction in ATP levels during ischaemia or pharmacologically by PIN, the increase in their open probability with stretch is potentiated. The additional stretch-induced changes in repolarisation (while potentially small, due to the transient nature of systolic stretch) may also be pro-arrhythmic, by further shortening APD and refractoriness, thereby increasing the period of excitability between normal beats. Here it should be noted that while APD_50_ can be used as an indicator of the transition from absolute to relative refractoriness, in ischaemia the recovery of excitability can lag behind repolarisation, a phenomenon known as post-repolarisation refractoriness.^65^ We did not quantify refractory periods in CTRL or SI using repolarisation-timed electrical stimulation protocols, but note that that premature contractions and sustained arrhythmic activity could be excited by late systolic stretch. This indicates that cardiomyocytes were excitable, and that the conceptual view that APD_50_ can be used to demark the start of a critical time window for mechano-arrhythmogenesis during RRC may be plausible. That said, it can be expected that the refractory period will differ between individual cardiomyocytes, which may be another reason why the overall incidence of mechano-arrhythmogenesis was not higher.

The finding that disturbed RRC, increased TRPA1 channel activity, and elevated [Ca^2+^]_i_ or ROS can lead to systolic mechano-arrhythmogenesis may have important implications for anti-arrhythmic treatments in various cardiac pathologies. In the case of myocardial ischaemia, previous experimental^12^ and computational^13^ studies suggested that ventricular arrhythmias originating at the ischaemic border are driven by systolic stretch-induced excitation and altered repolarisation, caused by mechano-sensitive ion channels interacting with ischaemic changes in the electrophysiological substrate. However, the molecular mechanisms responsible for this apparent systolic mechano-arrhythmogenesis were not experimentally investigated. As others have shown that during ischaemia: (i) RRC dynamics are altered;^14–16^ (ii) TRPA1 channels are activated;^56,57^ and (iii) [Ca^2+^]_i_ and ROS levels are increased,^11^ our results suggest that these various factors are mechanisms underlying systolic mechano-arrhythmogenesis in ischaemic myocardium.

Previous efforts to elucidate the presence and mechanisms of systolic mechano-arrhythmogenesis in the whole heart have been limited by the spatio-temporal complexity of cardiac electro-mechanical activity. Changes in the cardiac mechanical environment vary over a broad and heterogeneous range, relative to the timing of electrical cycles in individual cardiomyocytes across the heart. Our single cell experiments highlight that there is a critical window for mechanically-induced arrhythmogenesis during RRC. In the regionally ischaemic heart, we saw tissue stretch at the ischaemic border during mechanical systole, with the timing of its later portion – as well as stretch-release – coinciding with the (prolonged) period of RRC in that region. Other cases of end-systolic mechano-arrhythmogenesis – such as *Commotio cordis* (which can occur in healthy tissue)^66^ – also crucially depend on stretch-induced excitation of cardiomyocytes that have just regained excitability^36^ (with arrhythmia incidence in this setting reduced by block of K_ATP_ channels^67^, presumably due to a shift in the timing of RRC^36^). The same mechanism seems to be at play in the setting of long QT, in which repolarisation can outlast mechanical systole in regions of the heart, creating a negative ‘electro-mechanical window’ and a period during which local stretch of tissue that has just regained excitability by dyssynchronous contraction, rapid ventricular filling, or β-adrenergic stimulation-induced aftercontractions can trigger sustained ventricular arrhythmias.^68^ Also, in cardiac pathologies involving disturbed RRC, such as with β-adrenergic stimulation in heart failure^69^ or catecholaminergic polymorphic ventricular tachycardia,^70^ systolic stretch may underlie focal arrhythmogenesis due to the resulting increase in the duration of RRC.

An important consideration for the decrease in arrhythmia incidence seen in mechanically unloaded or non-contracting hearts in the present study is the associated change in metabolic demand. It has been shown that Langendorff-perfused rabbit hearts that are mechanically unloaded or non-contracting have a lower oxygen demand than those performing physiological work, resulting in a slower accumulation of NADH (an index of myocardial oxygen deficiency) and a slower and reduced shortening of APD.^71^ The difference in energy consumption of myocardium in physiologically-loaded *versus* mechanically unloaded or non-contracting hearts may account in part – along with reduced stretch of tissue at the ischaemic border – for the observed difference in arrhythmia incidence. Furthermore, as ion channel function is closely linked to the level of metabolic activity, decreasing metabolic demand may alleviate other pro-arrhythmic changes in cardiomyocyte electrophysiology,^72^ Even so, it should be noted that the pharmacological experiments demonstrating the importance of [Ca^2+^]_i_ for arrhythmogenesis in the whole heart were performed in contracting hearts with a physiologically loaded LV. Also, as optical mapping of APD was performed in non-contracting hearts, it is likely that we underestimated the degree of RRC prolongation during acute regional ischemia in the whole heart, which was about half of that seen in our isolated LV cardiomyocytes.

In conclusion, we have identified disturbed RRC and enhanced TRPA1 channel activity as molecular mechanisms contributing to stretch-induced changes in cardiac electrophysiology and a loss of protection against systolic mechano-arrhythmogenesis in acute myocardial ischaemia. Further, we have demonstrated that targeting mechanisms underlying altered RRC or TRPA1 activity, such as with local delivery of agonists or antagonists, may represent novel anti-arrhythmic strategies, with potential applications for other cardiac diseases involving RRC prolongation or an increase in the expression or activity of TRPA1.

## NONTSTANDARD ABBREVIATIONS AND ACRONYMS

AITC: allyl isothiocyanate
[Ca^2+^]_i_: free cytosolic Ca^2+^ concentration
CaTD: Ca^2+^ transient duration
AP: action potential
APD: action potential duration
CTRL: control
DPI: diphenyleneiodonium
K_ATP_: ATP-sensitive potassium channel
LV: left ventricle
NAC: N-acetyl-L-cysteine
NCX: sodium-calcium-exchanger
PIN: pinacidil
ROS: reactive oxygen species
RRC: repolarisation-relaxation coupling
SI: simulated ischaemia
TRPA1: transient receptor potential kinase ankyrin 1
V_m_: membrane potential

## STATEMENTS

## Acknowledgements

The authors thank Gentaro Iribe (Asahikawa Medical University, Asahikawa, Japan) and Keiko Kaihara (Okayama University, Okayama, Japan) for technical assistance with cell stretch. Carbon fibres were a gift from Jean-Yves LeGuennec.

## Sources of Funding

This work was supported by the Canadian Institutes of Health Research (MOP-342562, PJT-185904, and PJT-190009 to T.A.Q.); by the Natural Sciences and Engineering Research Council of Canada (RGPIN/04879-2016 and RGPIN/03150-2022 to T.A.Q.); by the Dalhousie Medical Research Foundation (Hoegg Graduate Studentship to B.A.C and Capital Equipment Grant to T.A.Q.); by the Canadian Foundation for Innovation (32962 to T.A.Q.); by the Heart and Stroke Foundation of Canada (G-22-0032127 and a National New Investigator award to T.A.Q.); by the German Research Foundation Collaborative Research Centre SFB1425 (DFG #422681845; speaker P.K.; members B.A.C., J.G., and R.P.); and by a personal Fellowship from the German Cardiac Society (to B.A.C.).

## Disclosures

None.

## Notes

### Competing Interest Statement

The authors have declared no competing interest.

